# Nanoscale Lubrication in Biosystems as Rationalized in Terms of Fractons and Spectral-Mechanical Properties of Networked Biopolymers in Ionic Solutions

**DOI:** 10.1101/2021.03.22.436506

**Authors:** A. Gadomski, P. Bełdowski

## Abstract

Articular cartilage is a natural tribochemical device just-designed by nature. Yet, a vivid debate goes on toward the mechanisms by which its nanoscopic viscoelastic properties facilitate lubrication in terms of ultralow static and kinetic friction coefficients. In this concisely conducted conceptual discussion, we wish to point out that a nanoscale tribomechanistic description based upon certain “viscoelastic quanta”, called fractons, expressing spectral-mechanical properties of viscoelastic nets under the influence of force/pressure factor(s), may contribute substantially to the elucidation of ultralow coefficients of friction in the articular cartilage of predictable relaxational response. Our example unveils a part of a mechanically responsive viscoelastic network, such as a tied piece of hyaluronan molecule, fit in an Edwards type tube, in which upon water–mediated interaction of lipids with the hyaluronan when subjected to loading at the nanoscale, consecutive stress-field and ion diffusion actions occur simultaneously. It results in a natural-logarithmic formula that interrelates a number of hyaluronan’s interactive residues, *N*, with certain molecular-elastic (an exponent *γ*) and surface-to-volume (nano-colloid type) characteristics of around 2/3 to emerge near thermodynamic equilibrium, that is to say after a frictional loading action performed. It enables to relate uniquely a value of the exponent 0 < *γ* < 1/2 with a virtual tribomicellization scenario of the nanoscale friction–lubrication event accompanied by inevitable tubular-milieu viscosity alterations at criticality when the quasi-static friction scenario shows up, preferably with *γ* → 1/3 from above for large enough *N* –s. A periodic vibrational super-biopolymer’s mode is exploited, leading to a change in the nanoscale friction-lubrication period from which an opportunity to involve an essential contribution to the (nanoscale) coefficient of friction arises.

**PACS numbers:** 71.10.+x, 81.30.Fb, 05.70.Fh, 05.60.+w

## 1. Introduction

Comprehending mechanisms responsible for the interfacial properties of articular cartilage (AC) surfaces at the nanoscale has long been sought after due to their formidable performance characteristics and because their dysfunction uncovers, for instance, the disease burden of conditions such as osteoarthritis.

In the recent research on ultralow friction conditions of natural articular devices, such as the articular cartilage appears to be, there is a quest for seeking reliable measures of coefficients of friction, and the measures of the friction-lubrication (and sometimes, of wear) efficiency, especially valid for the nanoscale [1].

Amongst many scenarios applied to versatile kinds of articular cartilages’ functioning [2], there appears a method, which is based on a scheme involving a dynamical system, characteristic of a certain production of biomolecular micellar aggregates capable of facilitating the intermediate-zone lubrication of two rubbing poroelastic natural surfaces immersed in the synovial fluid [3, 4]. Hyaluronic acid or hyaluronan (HA) is a long (of up to millions of Daltons in weight) linear polysaccharide comprised of alternating units of D-glucuronic acid and N-acetyl-D-glucosamine, which can be found directly at the surface of phospholipid membranes [5]. Its ability for physical cross-linking make it behave in a non-Newtonian way, transitioning from an elastic gel to a viscous liquid when placed under load. HA networks increase the vitreous humor’s viscosity, provide structural rigidity in complexes with aggrecan in cartilage, and contribute to lubrication in synovial fluid. Although its anomalous viscoelastic properties in bulk have long been known [6, 7], it is still not well understood how HA works alongside other molecules to produce low friction weight-bearing surfaces of joint cartilage. These surfaces are covered with multiple phospholipid layers (of a thickness up to 1500 nm), where the outermost layer comprises phospholipids, proteins, and proteoglycans. Therefore the interplay between phospholipids and hyaluronan can result in enhancing mechanical properties within natural joints.

The dynamical system proposed for facilitating the corresponding lubrication works in the boundary lubrication regime [2]. It consists of a dynamic interplay between two surficial phospholipid sub-fields, located either at the rubbing surfaces or out of them, i.e., in the middle-zone layer. They are, in general, coupled to one another (provided that a friction event proceeds) in a complex oscillatory manner, albeit in the simplified model [8], a linear coupling has already been assumed.

The friction-lubrication mesoscopic dynamical system introduced involves a tribopolymerization (or specifically, tribomicellization) kernel. This kernel renders a weak (viz linear) coupling with the surficial-phospholipid subfields (inter)action, and it is directly related to the kinetic/dynamic coefficient of friction. Since the kernel has been taken as (loading) time-dependent, it may if properly assumed, also contribute fairly to the mathematically legitimate closure of the overall dynamical system, very capable of unveiling the peculiarities of the articular-cartilage friction effect on the nanoscale-responsive biomaterial characteristics [2, 3, 4].

Which are the physical (nano-mechanical) factors that have made the said mathematical closure effective? The answer has got a “surprising super-structure”: These are the (discrete) interactive and innate nano-mechanical subfields that have arisen during a subtle nanoscale closure-making friction act. Namely, it is not doubtful that ultimately an “elementary” friction act of the two opposing soft-and-wet (nano)surfaces relies upon approaching (in a real mode) hyaluronan-decorated phospholipid nano-surfaces [2, 3] circumvented by the ionic milieus, presumably full of water and small ionic contents.

The latter supports very much our ionic-and-aqueous nano-surfaces approaching theoretical proposal, expressing the clue of our consideration for the present study [9]. The subtleties of the so-named proposal shall be disclosed below.

Thus, in the present study, substantial care has been taken to consider a biopolymer conjugate, made up of a linear hyaluronan chain as decorated with layered phospholipids, cf. Fig. 1, and co-supported by a necessary involvement of water dipoles assumed to be able to create *H*–bonds between the biopolymers and themselves.

**Fig. 1.**
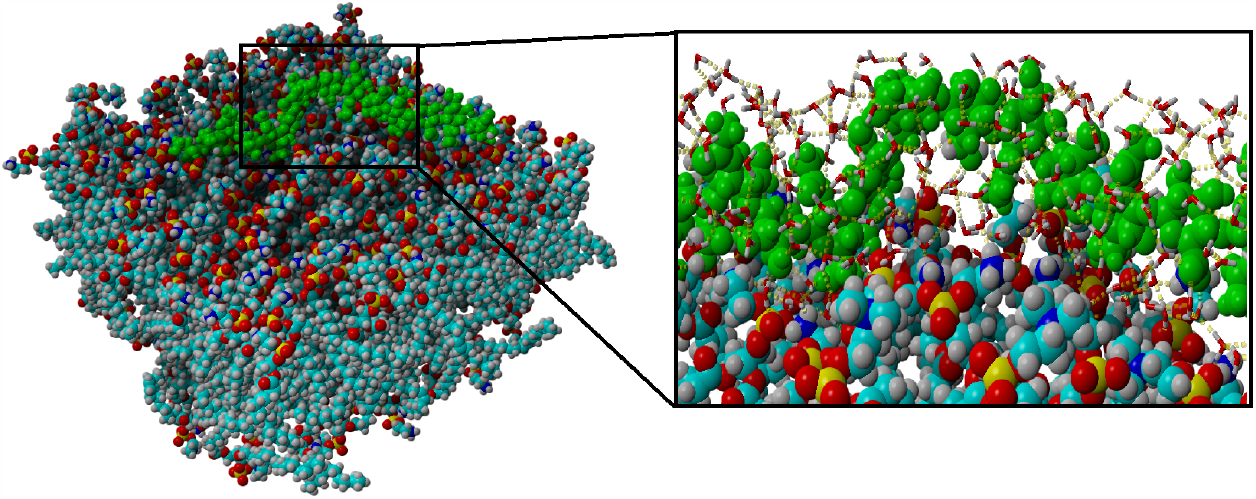
Snapshot of the final structure of the HA-membrane. HA is represented by the green colored chain. PL atoms are presented by colors as follows: oxygen-red, carbon-turquoise, hydrogen-white, phosphorus-yellow. Zoomed structure on the right-hand side shows a H-bond network of water-HA-membrane system. Hydrogen bonds are depicted as yellow dotted lines.

Because it is well-substantiated that hyaluronan can readily make the conjugates with other biopolymers, it is taken for granted that it renders a type of new biopolymer when decorated with the lipids. Since the conjugates of hyaluronan are mostly with proteins, in the current study, it is assumed that this hyaluronan-lipid (water assisted) conjugate, see Fig. 1, can be further used as water containing the proteinous equivalent of the biomolecular quasi-linear chain [6, 7].

When resting upon such assumption, we are fairly automatically encouraged to plunge into spectral-mechanical fractons addressing context, pointing to a quasi-equilibrium state of a super-biopolymer, composed of hyaluronan-lipid units immersed in an aqueous solution and having *N* active (lipids-holding) residues to which naturally a fractal geometry is suitably ascribed. Consequently, the number of residues (*N*) is, according to [10, 11, 12], related to the system’s spectral dimension. This quantity is, in turn, composed of the two geometrical dimensions: The fractal dimension of the “backbone” and its counterpart responsible for the liberated ions’ random walking along the “backbone”. The ions are generated by breaking the *H*–bonds when realizing the above-mentioned elementary friction act; these are preferably the hydrogen ions [3]. Of course, the friction act costs some mechanical energy to be dissipated at the nanoscale. It results in taking into account the actual mechanical state of the super-biopolymer, namely, whether it is - as a part of the hyaluronan’s net, and upon an elementary and effective load - a squeezed or a “normal” (viz Gaussian) chain, or it is an extended (swollen) chain [3, 4].

In this study, it is ascertained that a natural-logarithmic formula interrelating a number of hyaluronan’s interactive residues, *N*, with certain molecular-elastic (an exponent *γ*) and surface-to-volume (nano-colloid type) characteristics of around 2/3, emerging near thermodynamic equilibrium, unveils a most probable dynamic behavior of the super-biopolymer upon a static-friction event, that is, at the elementary frictional loading action performed. It then enables to relate in a unique albeit confined way a value of the exponent 0 < *γ* < 1/2 of the super-biopolymer with a tribomicellization scenario of the nanoscale friction–lubrication event accompanied by inevitable tubular-milieu viscosity alterations when the quasi-static friction scenario shows up, preferably with *γ* → 1/3 from above for large *N* –s. To make a long story short, such treatment leads to introducing in a fairly non-ambiguous way what types of kinetic friction coefficients are supposed to be predicted just in accordance with the restrictively preferred values of *γ*–s, as it has been discussed preliminarily in [3, 4]. From such estimates, one can also fairly well derive the static coefficient of friction as its kinetic counterpart’s maximum value.

In what follows, we shall first introduce the model, then give certain reliable, mainly computer simulation-based examples [10, 13, 14], and finally, discuss their reliability and robustness as well as an appreciable novelty of the rationale offered, especially when confronting it with certain previous studies, providing a nanoscale involving motivation to the current project [15], [16], [17], [18].

In the meanwhile, let us focus on the Grotthuss proposal that has provided a suitable conceptual framework to describe general prototropic transfer by hoping (i.e., by water-assisted proton involvement) from one water molecule to the next and associated with a suitable water molecule reorientation and mechanical agitation of the super-biopolymer’s vicinity [19, 20, 21]. The actual transfer mechanism, associated with the cascade of fractons and phonons, remains a subject of some dispute [15, 22], particularly because the identity of the conducted proton may change [23] and the recipient water molecule (attached to a lipid or a part of hyaluronan) may reorient in the hydronium donor field [24]. Water is a very suitable medium for proton conduction because of its propensity to form hydrogen bonds whereby the hydrogen-bonded network can delocalize solvated hydrogen ions. Hydrogen ions transport is determined by the rate at which hydrogen bonds between a hydronium ion and a water molecule form, which, when considering the hydrogen bond flicker rate of femto- to pico-second levels [25], is a very rapid process. It has further been proposed that proton/hydrogen ion mobility is higher than other ions because a certain contraction of the *O* − *O* bond distance adds energy to hydroxide transfer [26] and the additional kinetic energy of the excess proton augments the energy of hydronium ions [15].

## 2. Computer-Simulation Based Example

To demonstrate how it can be exemplified *in silico*, cf. Fig. 1, all-atom molecular dynamics simulations were performed using the AMBER14 force field [27] to evaluate interactions between one HA molecule and a dipalmitoylphosphatidylcholine, DPPC-DPPE (two most common lipids in synovial fluid [14]) bilayer. The HA structure was downloaded from PubChem [28] and modified to obtain longer chains using the YASARA Structure Software. The final molecular mass of HA considered in this study was 3 kDa. The DPPC-DPPE membrane consisting of 200 lipids was built with a ratio of 1:3, respectively. The bilayer was equilibrated for 10 ns, then HA was added to the surface and again equilibrated for another 10 ns. Periodic boundary conditions were applied to create an infinite bilayer. The TIP3P water model was used [29]. For the isobaric-isothermal ensemble, all-atom simulations were performed under the same conditions: temperature 310 K (physiological), pH=7.0 (acidic, reminiscent of man’s saliva). The time step was set to 2 fs, 0.9 NaCl solution. The radial distribution function (or RDF), *g*(*r*), is an example of a pair correlation function, which describes how, on average, the atoms in a (model) biopolymeric system are radially packed around each other. The radial distribution function (abbreviated by RDF), *g*(*r*), reads

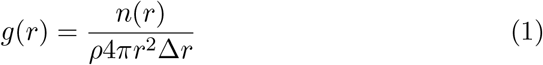

in which case *n*(*r*) is the mean number of atoms in a shell of width Δ*r* at distance *r* and *ρ* is the mean atom density. The method is not restricted to one atom - all atoms in the system can be considered, leading to an improved RDF determination as an average over many atoms.

Molecular dynamics simulations show the affinity between hyaluronic acid and phospholipid molecules (as presented in Fig. 1). The presented membrane comprises two phospholipid types with the highest abundance in synovial fluid: DPPC and DPPE. The interaction between HA and lipids is rather complex due to HA’s high hydrophilicity (see Fig. 2). Interaction between them considers three main interactions: hydrogen bonds, hydrophobic interactions, and water-mediated bonding. There is also another specific for lipid type. DPPC is more prone to form hydrophobic interactions and water-mediated contacts; DPPE forms more hydrogen bonds, i.e., creating more stable complexes. However, AC membranes are predominately composed of DPPC lipids, thus enabling HA along membranes [14]. HA interacts with phospholipids through its constituent polar (oxygen and nitrogen) and non-polar (hydrocarbon) parts. In Fig. 2, one can see RDF of water oxygen from all H-bonded HA’s atoms. This is useful to roughly determine the fractal dimension *d*_*f*_ from the following formula 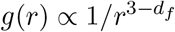 [30], where *d*_*f*_ is a fractal dimension. As one can see, RDF through the HA molecule changes depending on atom type. Carboxyl (O5 and O6) and hydroxyl (O1, O2, O8, and O10) group atoms show a high affinity to water, i.e., it can definitely participate in organizing lipids inside the membrane this way shaping its properties adequately [31]. Cell membranes show up a great fluidity, i.e., mobility of lipids inside - they are moving rather freely [32]. This, however, depends on lipid packing (a type of lipids inside the membrane). The membrane’s viscosity can affect the rotation and diffusion of proteins and other bio-molecules within the membrane, thereby affecting the functions. Moreover, lipids show up the inertial effects in the range of picoseconds, and a power-law decaying viscoelastic memory extends over the range of several nanoseconds [33]. This, in terms of HA-lipid binding, is related to the tribologically efficient mechanism of formation of inter-and intramolecular H-bonds, increasing the stiffness of HA (connected to the presence of water molecules that are increasing their persistence length) and stabilizing lipid membranes.

**Fig. 2.**
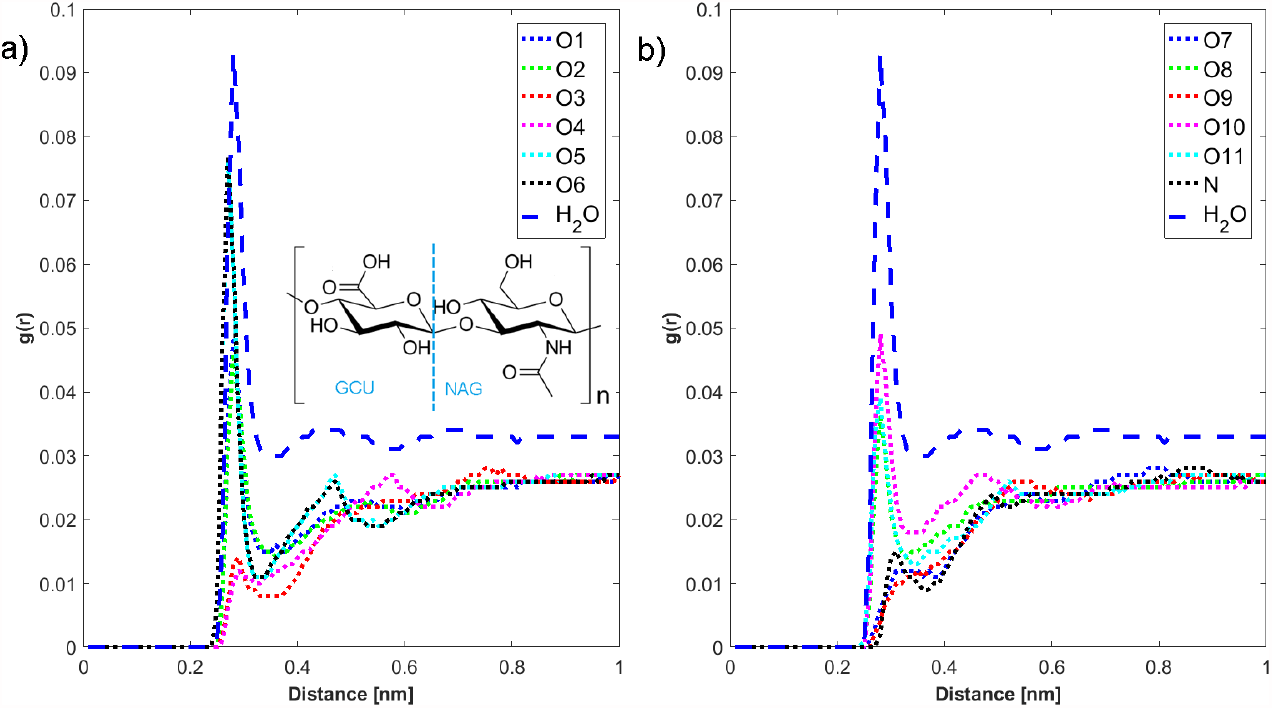
Radial distribution function for water oxygen atoms away from HA (structure shown). As one can see most oxygen atoms (carboxyl, hydroxyl group and O11) show similar behaviour i.e. fractal dimension remains constant throughout the polymer.

## 3. Fracton Type Model with Its Basics

The nanoscale-dynamics addressing model proposed in what follows is based on a vibrational mode that occurs when an elastic mesh in a HA network is slightly distorted from its natural (thermal) phonon mode. The distortion is often called a fracton mode, and it is realized, by contrast to the phonon low-frequency mode, in a higher frequency regime. This fracton mode is plausible to happen to the super-biopolymer, cf. Fig. 1, equivalent to the by-lipids-decorated hyaluronan chain embodied in an elastic mesh role of nanomechanical sensor after application of the corresponding load’s contribution transmitted to the nanoscale from its macroscale counterpart. Fig. 3 presents a coarse-grained representation of such a model as adopted from MD simulations.

**Fig. 3.**
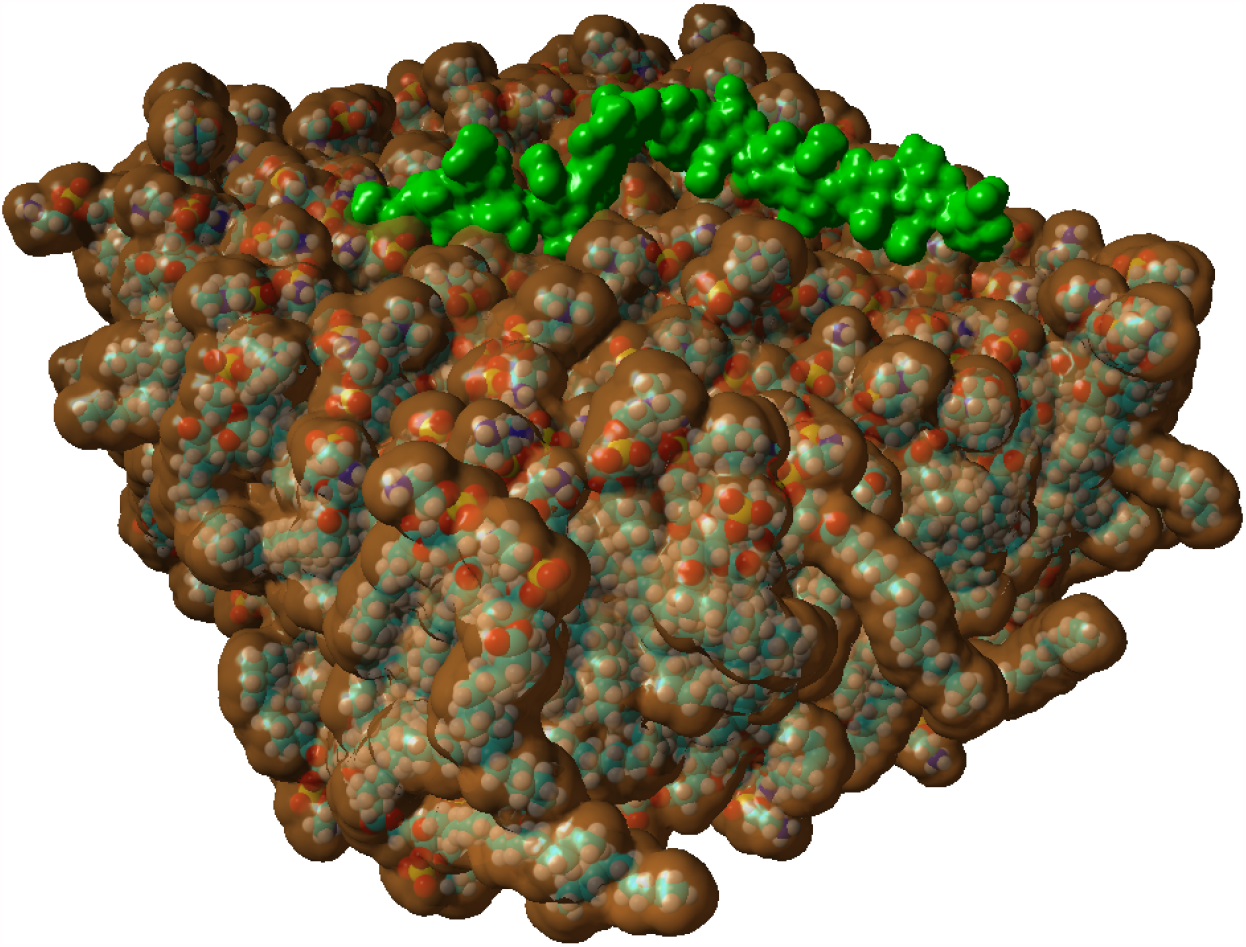
Coarse-grained representation of HA-membrane system. HA is represented by the green tube, whereas the phospholipid surface is represented as brown colored surface. Although length of HA used for MD simulations is much shorter than the one discussed in theoretical considerations, its interaction modes with phospholipids are the same as those of length of up to N=400 [31]. This, however, is also connected with lipid composition of a membrane/vesicle.

In several studies on nanoscopic friction-lubrication mechanisms [3, 4, 9] it was underlined that lipids, when associated to hyaluronan nanosurfaces, and immersed in an aqueous ionic solution, can provide a synergistic perspective on the very basics of nanoscale friction and lubrication. There can be three principal mechanistic modes that are very pertinent to the dynamics of three hydrogen-based accompanying ionic waves, propagating through the nanoscale parts (Fig. 3) of articular cartilage [4, 6].

In this brief presentation, we wish to confine ourselves to the parametric interval for which an elastic and characteristic exponent obeys 0 < *γ* < 1/2. What does it really mean? First, it means that we do not take into consideration all three nanolubrication modes mentioned [3]. Second, that we permit to introduce an after-load relaxation mode that relies on (a) compressing the molecules that would possibly (un)block the hydrogen ions passages; (b) after relaxing the molecular compression mode, one expects naturally to enter the so-called standard viz exponential, thus Debyean (consonant with diffusion), relaxation mode. The first squeezing mode leads to providing a maximal virtual space for the hydrogen ions to penetrate the pertinent articulation nanospace, whereas the corresponding second mode does prepare the nanoscopic part of the system to limit suitably the lubrication space. This type of confinement results in manifesting the lubrication space to disclose a type of random confinement with diffusive barriers that may freely show up in any articulation space [4, 10]. In fact, the physical situation is reminiscent of the one expressed by diffusion with the presence of entropic barriers as it has been described in [34].

When applying the macroscale load, it is accepted that a certain high-impact vibration can be transmitted into the nanoscale. In our opinion, this is the substantial case of the articular cartilage while functioning [2]. In fact, when there is no load transmission, the viscoelastic network under consideration fulfills the thermal phonon mode condition. If, in turn, an even small mechanical external force field applies, the vibrations lose their thermal-mode character, and they start to manifest readily with an increase of the expression frequency, thus entering their fracton or spectral-mechanical mode.

In the current study, it is proposed to rationalize the nanoscale friction-lubrication (FL) behavior by means of a harmonic spring, the characteristic period of which, denoted by *T*, changes within a time scale Δ*t* small compared to full-mode FL expression of the nano-spring, realized in time *t*, where Δ*t* << *t* holds. The full-mode FL expression is assumed to be equivalent to one spring cycle, occurring in *T*.

When embodied in a curvilinear or corrugated (circumventing) nanotube, the nano-spring, equipped with an aqueous solution (not shown in Fig. 3), consists of a piece of hyaluronan, decorated uniformly by layers of lipids. Of course, the so-invented nano-spring, as part of the viscoelastic mesh, has to be dealt with as exposed to either a horizontal or a vertical direction. These two directions are typically oriented either along with the load’s preferred orientation or perpendicularly to it.

In what follows, one is encouraged to assume an “average” mixing mode between the impacts caused by the two mutually perpendicular actions of the nanoload. It results in accepting an also averaged picture of the nano-spring’s dynamic behavior, given by

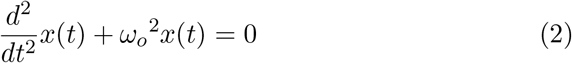

in which *x* represents a *t*-dependent distortion of the biomolecular spring from a local equilibrium position, and *ω*_*o*_ stands for the eigenfrequency of the ideal oscillator, a construct being equivalent to the super-biopolymer already introduced; cf. Fig. (3).

It is well-known that within the deterministic ideal-oscillator picture *ω*_*o*_ obeys *ω*_*o*_ = 2*π/T*, thus being inversely proportional to the characteristic oscillator’s period *T* [35, 36]. But one also knows that

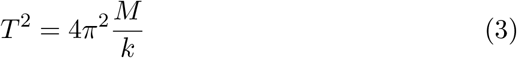

with *M* the total mass of the oscillating super-biopolymer, and with *k* designating its spring constant [37]. In a real FL (friction-lubrication) event with the eigenfrequency *ω*_*o*_ (or, *T*) it is legitimate to assign to it a stochastic picture based on the Poisson and/or birth-and-death mass-dependent process as it has been sketched in what follows.

Thus, it is natural to assume further [38] that the mass *M* of the oscillator undergoes a mass attachment-detachment effect on a rearranged decoration of the hyaluronan. Namely, if an elementary (force) nano-push will distort the corresponding viscoelastic mesh in which the chain is sitting, a number of lipids will detach from the hyaluronan’s surface [5] as to contribute to changing the viscosity of the surrounding (nanotubular) medium locally. The mass *M* would tend to decrease after the push. The spring, just in favor of its spring constant (*k*), in turn, have to relieve its bare (HA–characteristic) springiness, this way restoring its natural value, thus, providing a robust nano-spring without the lipidic overhang. Relieving its springiness means that the super-biopolymer will momentarily (in a small time interval Δ*t*) profit on its intrachain elastic-force contribution, having it -due to a loss of lipids’ mass - not engaged in the interchain (hyaluronan-lipid) associations, cf. Fig. 1. It is then plausible to take a Poissonian Δ*t*–effect [35] on *M* and *k* to be responsible for the further modeling in a simple linearized form in the Δ*t* interval. Thus, let us first propose

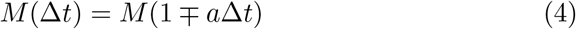

with *a >* 0 as an inversely time-dependent constant (in terms of physical dimension), pertinent to the detachment (interchain-interaction-weakening) kinetics introduced, thus pointing to a random subtraction from the total mass, see the sign preceding Δ*t* in Eq. (4), and

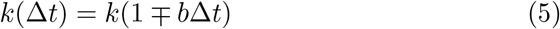

with *b >* 0, another (inversely time-dependent as well) constant, describing the intrachain interaction (randomly occurring) promotion, resulting in restoring momentarily the biopolymer/hyaluronan natural springiness (without lipids), because of the loss of lipid cluster mass got stuck to it before the friction event occured; cf. a very similar description of the Poisson process in [36]. Consequently, the rates *a* and *b*, as well as the signs preceding Δ*t* in both equations above, characterize how the super-biopolymer gains or loses on its overall intrachain interaction, making it either more elastic and solid or less elastically responsive, especially because of nanoscopic pseudoplastic or shear-thinning microrheological effects [2, 6], respectively. The first nanomechanical stage, devoid of the lipidic decoration, would correspond to the biopolymer as represented in terms of a Gaussian chain, thus exponent *γ* → 1/2. The second stage, as attributable to the FL mode for which the involvement of the lipid is crucial, would be ascribed to the case of the non-Gaussian viz compressed chain to be placed in the fractons’ regime with *γ* < 1/2, opting for *γ* → 1/3 [4]. As a consequence, one may expect a small non-perturbative modification in the time domain as follows

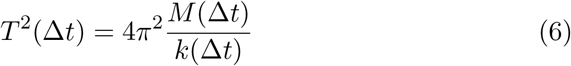

where Δ*t* << *t* applies. Put it in a simple explanatory manner, the attachment-detachment, super-biopolymer reconfigurable stochastic processes (characteristic of Δ*t*) [37], are typically much more quicker than their FL counterparts, proceeding in the friction-and-load interval *t*. This can be viewed as a type of adiabatic approximation characteristic of Δ*t* << *t*, leading typically to formal decoupling of the attachment-detachment and FL time scales. Bear in mind that the dynamical system that has already been presented for resolving a puzzle of the facilitated lubrication and friction at mesoscopic level of description [4], [8], was disclosed to be of inherently periodic character [3]. This periodic character is allowed to manifest in our modeling by including the reverse effect: The lipids go back to HA biomolecule (the plus sign in Eq. (4)), thus reconstituting the full super-biopolymer, whereas its springiness has to gain on its value because of having to carry the lipidic mass again (the plus sign in Eq. (5)). As argued below, it would correspond to a linear birth-and-death process.

## 4. Super-Biopolymer and its Friction-Lubrication/FL Properties

Given the afore presented, one can also say that there are two distinguished modes of the nanoscale friction-lubrication (FL) contribution to the overall articular-cartilage-based friction phenomenon. One of the modes for which *γ* → 1/2 represents the Gaussian super-biopolymer, whereas the second, presumably with *γ* → 1/3, represents the compressive non-Gaussian chain.

For the hyaluronan chain, in the context of the FL phenomenon to prevail, the number of its residues *N* ranges from 2.5 × 10^2^ to 2.5 × 10^4^ carbon (*C*_*α*_) atoms. (The values correspond to ca. (10^5^, 10^7^) Daltons, which is a realistic circumstance here.) For the case of the proteinous biopolymeric chain, a formula for the logarithm of *N* has been thoroughly checked numerically and improved based on two different fracton type (Landau-Peierls) models. It is proposed in a form of an “equation of biopolymer state” as follows [10, 11, 12]

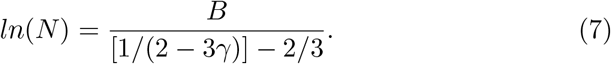

Notice that *B* is here a biopolymer type-dependent constant, most typically attaining *B* ≈ 4.5.

It is interesting if the above formula, working within the range of *N* –s from 2.5 × 10^2^ to 2.5 × 10^4^, coincides fairly well with the range of (1/2, 1/3). To give the *γ*–interval in more exact numerical terms, it results in having it as (0.44, 0.36(6)); thus, quite close to the fracton type model limits indicated elsewhere [3, 4]. (In ref. [3] a relationship between the decisive dimensions and exponent(s) has been established as 3*γ* + (*d*_*s*_/2) ≈ 2, with *d*_*s*_ being the spectral dimension, cf. Sec. 3 therein.) Especially, the limit *γ* → 1/3 from above, representing the fracton type FL mode, and pointing to the quasi-squeezed super-biopolymeric state, is worth appreciating here [3]. It is because this mode involves a bigger number of residues engaged in the FL event. Second, as described first in [3], it also allows a simultaneous action of the hydrogen ions, coming out from virtual breakages of the hydrogen bonds (Fig. 1) in the surrounding nanotubular milieu [9]. Third, it contributes to a structural rearrangement of the lipids; these biomolecules are assumed to rearrange into temporary nanostructures, facilitating the nanofrictional event to occur smoothly and predictably, cf. [2, 4]. Lastly, this can also be applied to understand a change in HA-phospholipids (mainly DPPC) interactions as dependent on HA molecular mass [31], cf. Eq. (7). HA of higher length (above 0,17 to 0,5 MDa) shows lower affinity towards lipid assemblies. High molecular mass HA shows higher persistence length thus resulting in lower binding affinity. Additional computer simulations (based on a coarse-grained approach), as well as experiments, could look into this specific problem to reveal the nature of so-envisaged phase transformation.

Such rearrangement already invoked may result in utilizing the adiabatic approximation (6) as embodied in the main oscillatory [38] super-biopolymeric mode governed by (2) and supported by the simplified Poissonian *M* vs. *k* temporal (counting) statistics, extendable to their birth-and-death counterparts, i.e. (4) and (5), respectively. The adiabatic or time-decoupling approximation does not take into account the critical conditions in which the periodic-in-nature friction act shows up. The criticality, expressed either by pseudoplasticity- or roughness-induced nano-effects (super-biopolymer vs. asperity interaction) to be switched on, can make a circumstance [14], resulting in a kinetic-friction hindrance, and causing the system to attaining a forced equilibrium value to be surmountable after a certain time *t* ≃ *T*.

As one is aware of mathematical involvement of time-variable *t* in Eq. (2), one agrees that a condition of semi-infinite axis of real values may be well assigned to *t*, namely that *t* ∈ [0, ∞) holds [16, 17, 39, 40]. In almost all circumstances, namely, excluding the ones at the afore described criticality, the adiabatic approximation prevailed, albeit it did not exclude the limit of Δ*t* →∞ to be applicable for the full FL mode, that means, the one having included the criticality circumstance. The large Δ*t* limit would favor the attachment-detachment kinetics to become readily operational because of its manifestation in all circumstances of the FL event, within the time domain reserved for the full friction condition to apply as its static-friction part, the part coinciding well with a mechanistic quasi-equilibrium condition is going to show up.

As a consequence, it is postulated to allow to make an exlusion from the almost-overall time scale decoupling by introducing the mode-coupling circumstance at criticality, and then by applying formally the L’Hospital rule, thus performing the limit of Δ*t* → ∞ on (6) which yields

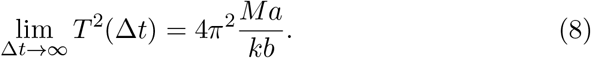

It is equivalent, after invoking (3), that

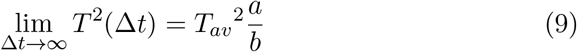

actually applies, albeit from now on *T*_*av*_^2^ = 4*π*^2^ < *M/k >* is an averaged value, and a crucial modification of the oscillation period appears to be the case. The modification is represented by the quotient *a/b* that stands for a bare interplay between two decisive sub-modes of the nanoscale FL mode. It is that the *a* parameter shows up the elastic intramolecular interactions addressing submode while the *b* parameter reveals its viscous super-biopolymer’s nearby milieu involving counterpart. The quotient *a/b* is a ratio between intra- and intermolecular interactions as applied to the super-biopolymer under a nano-load, causing the corresponding viscoelastic-mesh distortion by *x*(*t* ≃ Δ*t*) to show up in full, plausibly also in terms of a slight shear-thinning/pseudoplastic or super-biopolymer vs. asperity interaction co-effects to manifest efficiently.

The latter scenario disclosed, permits to try out a measure of the static coefficient of friction (COF) directly as *a/b*, wherein *a* and *b* are positively defined parameters; when designating it by *f*_*eq*_ one arrives at

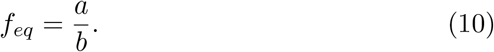

The kinetic COF, in turn, as denoted by *f*_*kin*_, would be then defined as

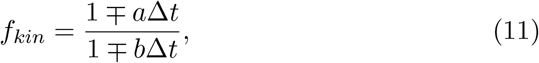

as it typically occurred for any of articular cartilages, having ultralow COF values [2]. In a typical FL circumstance it means that *f*_*kin*_ is going to be a weakly loading-time dependent observable that for the restriction of *a* << *b* (and, the postulated inclusion of the coupling time mode at criticality) behaves as *f*_*kin*_ ∼ *a/b*[1 ∓ (1*/a*Δ*t*)] at *t* ≃ Δ*t*. If *a*Δ*t* is big enough, its reciprocal value is appreciably small, thus, small compared to one, just arriving at the limit of static COF here, *f*_*eq*_ = *a/b*, see Eq. (10).

Based on the criticality involving conjecture (10), and after invoking (9), one has to provide

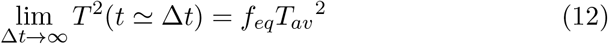

for a suitable experimental check-up. The check-up should take into account a periodic and critical loading process in the nanoscale (as applied to the nanomesh), with the propagation of the after-load stress over an effective length (*N*) of the hyaluronan [5] as (un)decorated with lipids, with the total (un)loading action to be detected within the nearby super-biopolymer’s nanotubular, curvilinear surroundings, equipped with ionic aqueous solution [2, 22, 26]. (The check-up of subtle nature can be reasonably thought of as realizable when employing tapping-mode AFM technique.) The full period *T* lasted over a swollen state of the super-biopolymer, with *γ >* 1/2 [3], would not be taken into account for the present modeling, then the propagation of the nano-load over superbiopolymer’s chain should proceed, starting from its quasi-equilibrium, Gaussian chain state with *γ* ≈ 1/2, thus upon the corresponding recovery of lost lipidic mass, and finally, upon landing on the squeezed (*γ* ≈ 1/3)–state, caused by another friction event, slightly out of equilibrium. In this sense, the friction event of interest, going forth and back in the course mass-springiness critical recovery, resembles a birth-and-death process studied in [36], having a randomly occurring diffusion of mass. It also reveals its inherent periodic character as argued in [2, 8], a physical fact being fully consistent with the proposed model. The method proposed for the experimental check-up is reminiscent of the (quartz) microbalance lubrication measurements, see [41]. For immersion in a suitable dynamic channeling context, one is encouraged to get in [43, 44].

Referring to the nanoscale and microbalance type measures explored above, let us evaluate a difference between (3) and (12), denoted by Δ*T* ^2^_*F L*_. It is easily calculated to be

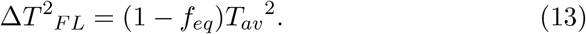

It tells us that the smaller the value of *f*_*eq*_ can be (0 < *f*_*eq*_ << 1), the more ‘ideally harmonic’ the FL elementary event is expected to be. Thus, it is profitable to work with super-biopolymers such as the ones exemplified by those included in the Introduction. A more advanced scenario of the FL process can be drawn in terms of the stochastic ideal-harmonic terms, as addressed by [35, 39]. As a consequence, *a* and *b* ought to be dealt with as stochastic-kinetic coefficients, in the most probable case arriving at their typical values, possibly within the bounds of *a* << *b*.

## 5. Summary

The current study’s main motivation was to find a reliable passage from macroscale to nanoscale within the realm of a friction-lubrication (FL) phenomenon pertinent to the functioning of AC systems (of *f*_*eq*_–s as low as 10^−4^) [17], thus, unveiling peculiarities of the action of incommensurable rubbing nanosurfaces composed of super-biopolymers. Conveying the model’s conception, it was assumed that when no sign of a macro-load has been transmitted to the nanoscale, the systems sit in a low-frequency (thermal) regime. If, in turn, the mechanical transmission mentioned becomes effective, it then causes certain changes to the super-biopolymer’s (Fig. 1 and Fig. 3) conformation. Consequently, a high-frequency fracton mode emerges, leading to a reconfiguration of the super-biopolymer’s structure. This event can be repeated after the next friction nano-event will become effective. After removing the cause of friction, the system relaxes to its local equilibrium state.

Such a conceptual proposal leads to the following striking dependencies between the observables responsible for a periodic behavior of the FL viscoelastic and nanoscopic system. These are:

i. First, the harmonic approximation to the FL system has been proposed in its adiabatic mode, pointing to the invariance of a ratio between its total energy (Hamiltonian) and its eigenfrequency [40], Section 3.
ii. Second, it turns out that the so-posed harmonic approximation can operate within the realm of a respective eigenfrequency (or period) modification that takes readily into account the corresponding structural changes of the super-biopolymer’s structure, viewed in terms of lipids’ detachment and by augmenting the elastic modulus of HA.
iii. Third, because the period’s modification in its magnitude is also innate to the full FL event, one is capable, after including the criticality conditions, of extracting the static and kinetic COF, just from the two stochastic-kinetic and structural co-events responsible for the overall effect on the FL system, thus, from making use of appropriate estimations of *a* and *b*.
iv. Fourth, as a consequence, such a simple procedure of extracting the COF–s enables one to get the values of COF–s in terms of the ratio between two main parameters responsible for the two concurrent and criticality addressing effects introduced.
v. Fifth, provided that we are speaking about a normally responsive AC system [2], the COF would be of proper very low value, such as *f*_*eq*_ ∼ 10^−4^, if the elastic contribution of HA (in fact, a rate of relieving the HA springiness) will win over its lipids’ detachment addressing counterpart, see Section 4. This can also be used to comprehend a change in HA-phospholipids (mainly DPPC) interactions as dependent on HA molecular mass [31]. HA of higher length (above 0,17 to 0,5 MDa) exhibits lower affinity towards lipid assemblies. High molecular mass HA shows higher persistence length thus resulting in lower binding affinity. Additional computer simulations (based on a coarse-grained approach), as well as experiments, could look into this specific problem to reveal the nature of above mentioned phase change.
vi. Sixth, as for the fracton characteristics involved in the FL elementary phenomenon just disclosed, one can employ in reality the RDF tool (Fig. 1 and Fig. 2; cf. Sec. 1 and Sec. 2) as well as the conception of super-biopolymer’s equation of state uncovered in Sec. 2, Sec. 3 and Sec. 4, involving the two important limits for the fracton-mechanical [12, 11] exponent *γ*, namely *γ* → 1/3 and *γ* → 1/2 [9].
vii. Seventh, the entire framework introduced in Sections 2–4 renders the system ready to examine particular interesting examples from computer simulations and experiments. The examination can then guide specific friction-lubrication scenarios occurring in healthy [42] as well as osteoarthritic articulating systems of interest [14, 2, 17], also leading to (not revealed explicitly) ion-channeling co-effects associated with the FL elementary phenomenon, cf. [3, 4, 9, 43]. For example, when commencing with another future study, it is to think in a legitimate way of exploring an FL stochastic extension based on approximate solutions to the Kuramoto model, characteristic of synchronized channels’ dynamics [44].

## Acknowledgements

The paper is to commemorate a great researcher/physicist, Professor Lutz Schimansky-Geier (1950-2020) of Humboldt University in Berlin, who passed away in Greifswald on November 2020. The work is supported by BN-10/19 of the Institute of Mathematics and Physics of the UTP University of Science and Technology in Bydgoszcz.

